# Repurposed small molecule toxin inhibitors neutralise a diversity of venoms from the Neotropical viperid snake genus *Bothrops*

**DOI:** 10.64898/2026.01.02.697350

**Authors:** Rachel H. Clare, Adam Westhorpe, Emma Stars, Taline D. Kazandjian, Laura-Oana Albulescu, Stefanie K. Menzies, Nicholas R. Casewell

## Abstract

Snakebite globally claims more than 100,000 lives per year and results in morbidity for 400,000 survivors. Current treatment uses antibody-based antivenoms which are constrained by their efficacy, safety and cost. In this study we evaluated the efficacy of previously described repurposed drugs against viperid snakes of the medically important *Bothrops* genus. Despite variable toxin representation and bioactivity across this central and south American genus, we found that the lead inhibitors targeting metalloproteinases (marimastat and DMPS) and phospholipases (varespladib), demonstrated pan-species neutralisation in enzymatic assays, whilst nafamostat (serine protease inhibitor) had variable activity. The metalloproteinase inhibitors protected against the procoagulant and haemorrhagic effects of several venoms in phenotypic assays. Collectively these findings demonstrate that repurposed drugs may be of great value as early interventions for the treatment of bothropic envenoming in the Neotropics and thus provides a strong rationale for their progression into future preclinical and clinical evaluation for snakebite indication.

## Introduction

Snakebite envenoming is an acute, potentially lethal event that affects several million people each year, primarily those working and living in rural areas of the world’s tropical and sub-tropical regions. Snakebite is classified by the World Health Organization as a priority neglected tropical disease, and is responsible for over 100,000 deaths annually, while several times that number of survivors suffer from long-term disabling or debilitating health conditions caused by venomous snakebites ^1^. Current treatment for snakebite consists of intravenously delivered antivenoms, which are serotherapies consisting of polyclonal antibodies generated from venom-immunised animals (e.g. equines, ovines, camelids) ^2,3^. Multiple venoms are often used in the immunising mixture, resulting in a broad range of antibodies that target the diversity of toxins present. Nonetheless, antivenom therapies are generally constrained in their efficacy to particular geographical regions or snake species due to venom toxin variation, which is highly variable across medically important venomous snakes ^4^.

The cocktail of toxins produced in the venom gland varies not only across snake species, but can also vary between populations and individuals of the same species, and has been shown to be affected by the age, sex and ecological factors experienced by the producing animal ^5–8^. The application of ‘venomics’ approaches over the past two decades has enhanced the identification and characterisation of snake venom toxin diversity at the proteomic level ^9^. This advancement, alongside functional characterisation, has supported the consensus that certain venom toxin families are often the dominant drivers of pathology following snakebite, due to both their high abundance and toxicity; namely, the snake venom metalloproteinases (SVMP), phospholipases A2 (PLA_2_), snake venom serine proteases (SVSP) and three finger toxins (3FTx) ^10^. This knowledge has paved the way for the discovery and development of alternative therapeutic approaches to antivenom, focusing on the development of rational data-informed therapeutic molecules which inhibit specific venom toxin families or sub-families, unlike conventional antivenom. Recent advances ^11^ include the identification of toxin-specific monoclonal antibodies ^12^ or nanobodies ^13^, computationally designed binding proteins ^13,14^ and synthetic inhibitors ^2,10,15^. Repurposed drugs (synthetic inhibitors), defined by the new therapeutic use for existing molecules previously developed for other indications, are showing considerable promise for snakebite envenoming. This is partly because three of the four priority venom toxin families to neutralise are enzymes (PLA_2_, SVMP and SVSP), lending themselves to potential generic toxin family inhibition (i.e. across diverse isoforms) by small molecule drugs via binding to the active site. Several such drugs have been shown to rescue the lethal and pathological effects of snake venoms in small animal models, including SVMP-induced coagulopathy and haemorrhage, PLA_2_-mediated neurotoxicity, and localised tissue damage via cytotoxicity (caused by PLA_2_ and SVMP) ^15–19^.

Several of these repurposed drugs have now entered or are entering clinical trials as orally bioavailable therapeutics amenable for clinical evaluation as front-line snakebite treatments. For example, the PLA_2_ inhibitor varespladib recently completed a Phase II trial for snakebite in the USA and India ^20,21^, with a second Phase II study ongoing at the time of writing.

Similarly, the SVMP inhibiting metal chelator DMPS has undergone Phase I studies ^22^ to dose optimise for snakebite indication, and is planned to enter Phase II evaluations in the near future alongside another SVMP inhibitor, marimastat, which has a distinct mode of action. Potential key benefits of such small molecules over traditional polyclonal antibody-based antivenoms include clearly established and acceptable safety profiles, broad-spectrum toxin inhibition, low-cost manufacture, improved tissue penetration and oral bioavailability ^15^. The latter provides a potential paradigm shift for treatment, from: i) the requirement for intravenous delivery (in tertiary healthcare settings) of antivenom which leads to treatment delays, ii) the reliance on cold chain storage and iii) the risk of serious adverse events from animal-derived antibodies, resulting in a vision of community based, rapid delivery of low cost, well tolerated, temperature stable, oral medications. Substantially reducing the time frame between bite and treatment, which in many settings such as the remote areas of the Amazon has been reported to be more than five hours, holds much promise for improving patient outcomes for this time sensitive, acute, life-threatening condition ^23,24^.

In this study we focused on assessing the potential utility of small molecule drugs in the context of the Neotropics. Unlike many other locations at risk of snakebite where a range of genera are indicated in the majority of severe envenoming cases, medically important snakebites in Central and South America are often dominated by snakes of the genus *Bothrops* ^25–28^. This group of diverse pit vipers (family Viperidae) includes species that range geographically from Mexico to Argentina, as well as certain Caribbean islands (Figure 1A). Conservative estimates of snakebite incidence in Latin America were previously reported as 50.37/100,000 people per year (80,329 snakebites per year), resulting in 540 deaths per year (0.5781/100,000 per year) ^29^. A more recent publication focusing on South America reported the continent as experiencing the third highest incidence of snakebite after Asia and Africa at 21.7/100,000 population per year, resulting in the fourth highest mortality rate at 0.03/100,000 population per year, after Asia, Africa and North America ^30^. The fact that Costa Rica and Nicaragua are classified within the North American dataset further exacerbates the snakebite incidence in the wider biogeographical realm of the Neotropics.

**Figure 1.**
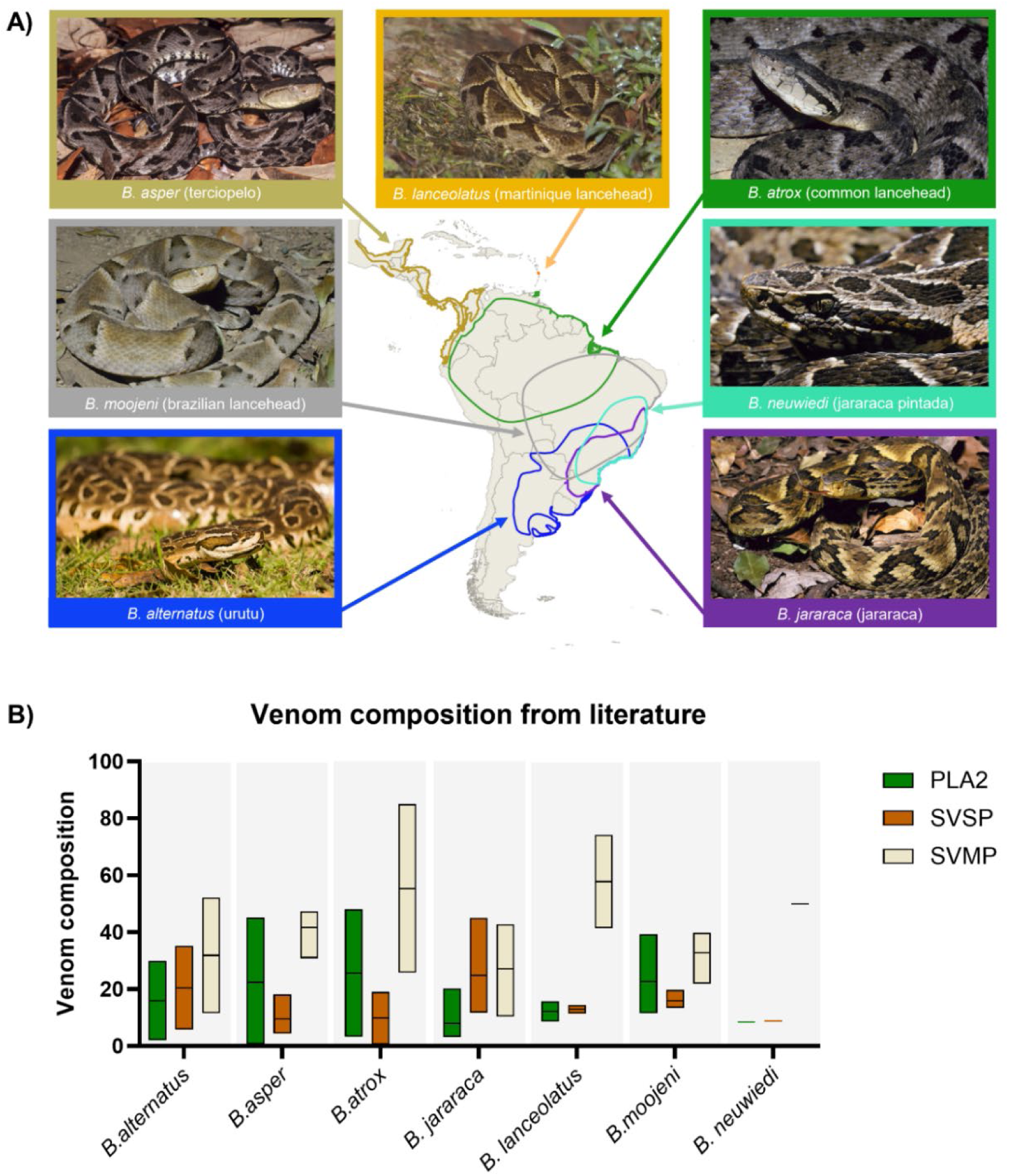
Variation in the distribution and venom composition of *Bothrops* species. A) Geographical distribution of the *Bothrops* species relevant to this manuscript. The distribution is based on the ICUN Red List accessed in April 2025 presented using QGIS 3.4. The images of *B. asper*, *B. atrox*, *B. jararaca, B. moojeni* and *B. neuwiedi* are held under all rights reserved copyright and have been published with permission by Wolfgang Wüster, while the image of *B. lanceolatus* is published with permission by Jonathan Florentin. The image of *B. alternatus* was accessed via Wikimedia commons and is held respectively under Creative Commons Attribution-Share Alike 2.0 Generic licence by Cláudio Timm. B) A summary of the published percentage venom compositions for the three key toxins of relevance to this manuscript; snake venom metalloproteinases (SVMP) in beige, snake venom serine proteases (SVSP) in orange, and snake venom phospholipase A2 (PLA_2_) in green. The extracted data (Supplementary Table 1) vary from an individual to large pools of specimens including captive and wild caught. The data is summarised from the following number of publications per species; *B. alternatus* (n=2), *B. asper* (n=2), *B. atrox* (n=6), *B. jararaca* (n=3), *B. lanceolatus* (n=2), *B. moojeni* (n=2) *and B. neuwiedi* (n=1). The data is displayed as a box and violin plot (interleaved low-high with line at the mean) using Prism software version 11 (GraphPad).

There are several commercial antivenom options to treat bothropic envenoming in the region, including the polyvalent antibotrópico antivenom from Instituto Butantan (Brazil) and PoliVal-ICP from Instituto Clodomiro Picado (Costa Rica). However, each antivenom is designed to cover only certain species, and delays between bite and accessing antivenom treatment remain, with a clinical epidemiological study of *Bothrops* bites in Brazil ^24^ estimating an average of three hours between accident and initial medical care for 65% of cases, but increasing to up to six hours for most cases in northern and northeastern regions of the country ^24^. Indeed, antivenom distribution centres are disproportionately spatially distributed compared to snakebite incidents, with the northern regions that experience higher incidences of *Bothrops* envenoming having fewer centres ^24^. These findings were mirrored in a systematic review of antivenom use in the Americas, which identified an average of 5.7 hours between bite and antivenom administration, with the treatment time ranging from 1.5 to 19 hours ^28^.

The typical clinical presentation of ‘bothropic syndrome’ following *Bothrops* envenoming includes local tissue damage (bruising, blistering, dermonecrosis, myonecrosis, and oedema), pain, incoagulable state, haemorrhage, circulatory shock and acute kidney injury. The latter three cause the most serious systemic and life-threatening effects ^24–26,31–33^. The reported pathologies linked to *Bothrops* envenoming are attributed to the dominant venom protein families, namely the PLA_2_, SVMP and SVSP enzymes (Figure 1B). To investigate the potential utility and therapeutic value of small molecule drugs for bothropic envenoming, in this study we tested the ability of previously described repurposed drugs to neutralise a wide range of *Bothrops* venoms in a panel of toxin-specific enzymatic assays, followed by phenotypic assays measuring venom-induced coagulopathy and general haemotoxicity. Our findings demonstrate that the lead repurposed snakebite drugs currently under study (varespladib, marimastat and DMPS) provide broad, albeit often variable, inhibitory potency against the diverse enzymatic activities of *Bothrops* venoms. Furthermore, the SVMP inhibitors show much promise for inhibiting the coagulopathic and haemotoxic effects caused by these venoms, providing a strong rationale for their future preclinical and clinical evaluation.

## Results

### *In vitro* profiling of *Bothrops* venom activity

#### Variation in venom composition

Figure 1B summarises the published reports of the venom toxin compositions of the species tested in this manuscript. These prior studies demonstrate clear interspecies variation in venom toxin composition, but also intraspecies variation for several cases (Supplemental Table 1). In this manuscript we did not seek to formally characterise venom composition, but instead used SDS-PAGE to provide a contextual overview of the venom protein compositions of the species used in this study (Figure 2A). In line with the literature, notable differences were observed between the species in terms of the proportion and abundance of proteins of all molecular weights. The seven venoms could be broadly assigned into two groups. The first group, containing *B. alternatus*, *B. jararaca* and *B. lanceolatus* venoms, exhibited abundant protein bands at ∼55-60 kDa (likely PIII SVMPs) alongside less abundant lower molecular mass proteins at ∼20-35 kDa (likely SVSP, cysteine-rich secretory proteins and/or PI SVMP) and ∼12-16 kDa (likely PLA_2_ and C-type lectin-like proteins) ^34^. The sample of *B. jararaca* used in this study had a dominant protein band ∼55-60 kDa, which if assumed to be PIII SVMPs, was in line with previous proteomic studies (Figure 1B, Supplementary Table 1) ^35^. The second group, containing *B. moojeni*, *B. neuwiedi*, *B. asper* and *B. atrox* venoms, had a much higher abundance of lower molecular weight toxins (<25 kDa), including prominent bands at ∼12-16 kDa, and lower abundance of higher molecular weight toxins.

**Figure 2.**
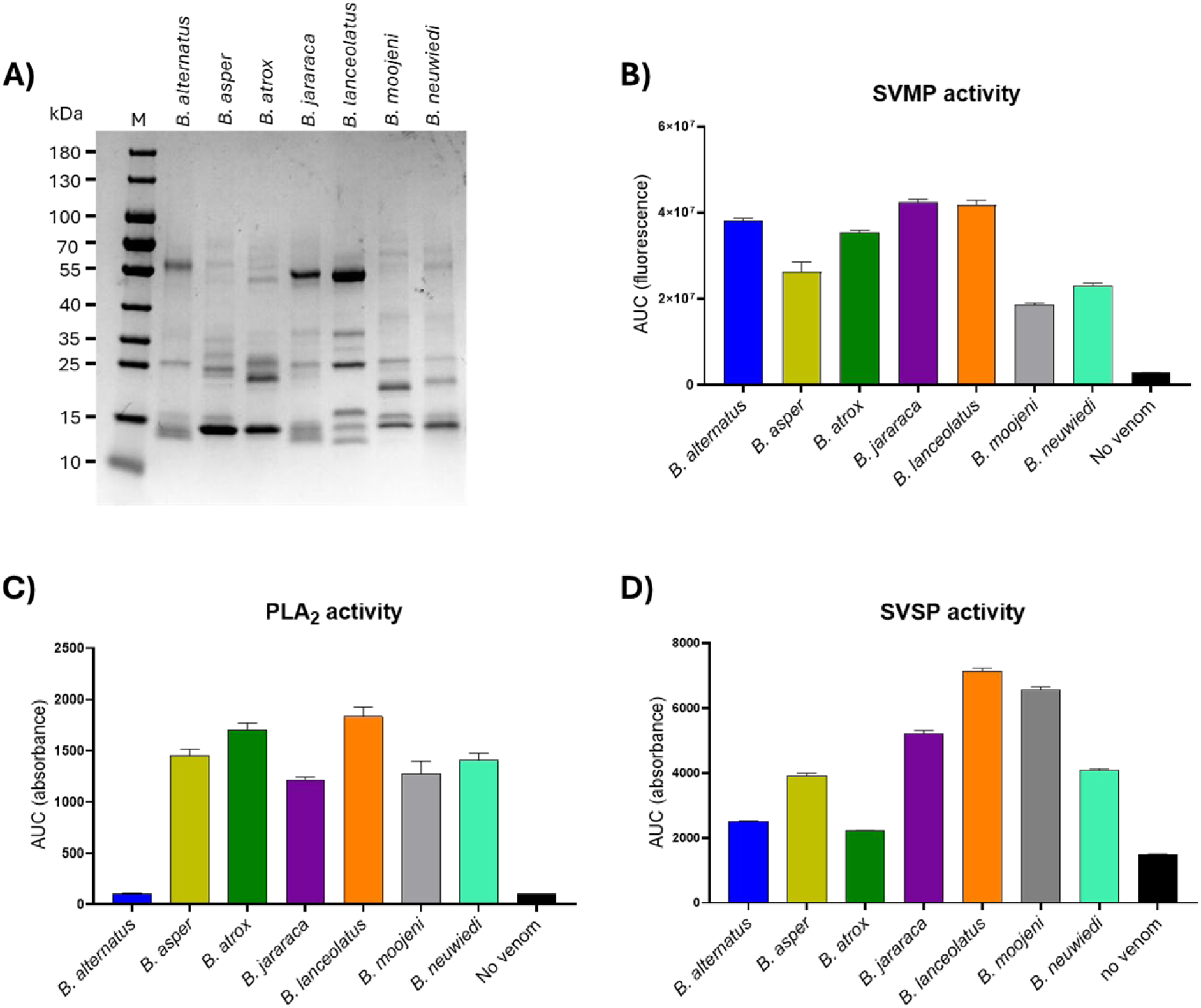
*In vitro* profiling of venom samples from seven different *Bothrops* species. A) Protein profiles of the venom protein components per species. Whole venom (5 µg per lane) was prepared under denaturing and reducing conditions, then separated by SDS-PAGE using a 4-20% gel. The gel was then stained with Coomassie blue and destained to show all proteinaceous components. M = molecular weight marker, and approximate masses of the markers are given in kDa. B-D) enzymatic activity of the seven *Bothrops* species in toxin specific assays. The results were analysed using area under the curve (AUC) in seconds and plotted using Prism version 11 software (GraphPad). A) Snake venom metalloproteinase (SVMP) activity using fluorogenic substrate ES010 read at an excitation wavelength of 320 nm and an emission wavelength of 405 nm over 108 minutes (n=3, ±SE), with 1 µg of venom per reaction. C) phospholipase A_2_ (PLA_2_) activity using commercially available secretory PLA_2_ kit (Abcam) read at an absorbance wavelength of 405 nm over 12 minutes (n= ≥2, ±SE), with 100 ng of venom per reaction. D) snake venom serine protease (SVSP) activity using commercially available chromogenic substrate S-2288 read at an absorbance of 405 nm over 35 minutes (n=4 ±SE) with 1 µg of venom per reaction.

#### Comparisons of SVMP activity

Quantification of venom SVMP activity used a consistent venom and substrate concentration and was measured kinetically. The area under the curve (AUC) of each kinetic reaction was calculated, and this value was used to compare SVMP activity between the venoms (Figure 2B). All venoms (1 µg per reaction) demonstrated some degree of SVMP activity; *B. jararaca* and *B. lanceolatus* displayed the most activity (AUC = 4.24 × 10^7^ and 4.18 × 10^7^ respectively) followed by *B. alternatus* (3.82 × 10^7^) and *B. atrox* (3.55 × 10^7^), then a moderate drop in SVMP activity of 1.4 to 2.3-fold was observed with the final three venoms (*B. asper*, *B. neuwiedi* and *B. moojeni*; 2.62 × 10^7^, 2.31 × 10^7^ and 1.86 × 10^7^, respectively). Only the four most active venoms had converted all the substrate present in the reaction by the end of the experiment, causing a plateau in fluorescent signal. A general trend was seen between the total intensity of the higher molecular weight SDS-PAGE bands that corresponded with PIII SVMP proteins (i.e. ∼55-60 kDa in Figure 2A) and SVMP assay activity, with the exception of *B. atrox*, however the potential contribution of other SVMP subclasses to this activity remains unclear.

#### Comparison of PLA_2_ activity

Venom activity in the PLA_2_ assay also showed considerable variation. As with the SVMP assay, the AUC of the curve generated from the kinetic read of the assay was used to compare PLA_2_ specific substrate conversion rate, and therefore PLA_2_ activity, of the venoms tested at a matched concentration of 100 ng per reaction (Figure 2C). *B. lanceolatus* and *B. atrox* venom demonstrated very similar and potent PLA_2_ activities (AUC = 1832.0 and 1703.0, respectively), with activity 1.2 to 1.5-fold higher than the activity seen to the next grouping of venoms, which all displayed very similar activities; *B. asper* (1449.8), *B. neuwiedi* (1408.5), *B. moojeni* (1269.5), and *B. jararaca* (1214.5). *B. alternatus* displayed no PLA_2_ activity at this dose (105.2 compared to the 99.56 assay background) and was distinctly lower than any of the other tested venoms, but this correlates with the previously described venom composition of *B. alternatus* (2% PLA_2_; Supplementary Table 1) ^35^. Correlations with the SDS-PAGE in the absence of proteomic data are challenging, as all venoms showed protein bands that corresponded with the predicted molecular weight of PLA_2_ toxins (∼12-14 kDa; though these also overlap with other venom toxins), including *B. alternatus*, though the strongest bands were observed in *B. atrox* and *B. asper* venom which were the second and third most active in this enzymatic activity assay. While *B. lanceolatus* showed bands of weaker intensity at this molecular weight, its venom profile showed three distinct bands in this molecular region, which may cumulatively account for the highest functional activity seen across all of the venoms tested.

#### Comparison of SVSP activity

Quantification of the kinetic profiles of SVSP activity revealed more extensive variation compared with the findings from the PLA_2_ and SVMP activity assays. The venoms of *B. atrox* and *B. alternatus* presented with low levels of enzymatic SVSP activity (mean AUCs: 2213 and 2504), compared to the moderate activity of *B. asper*, *B. neuwiedi* and *B. jararaca* (mean AUCs: 3916, 4073 and 5214) (Figure 2D). The highest venom activity was observed for *B. moojeni* and *B. lanceolatus* (mean AUCs: 6552 and 7143), which exhibited greater than 2.5-fold higher activity than *B. alternatus and B. atrox* venoms and between 1.2 and 1.8-fold than that of the moderately active venoms of *B. asper*, *B. neuwiedi* and *B. jararaca*.

#### Comparison of coagulation profiles

The plasma coagulation assay was conducted in a similar manner to the previous assays, in which a fixed venom dose (100 ng per reaction) was used to compare the coagulopathic activities of each venom. This assay utilises a biological product, citrated bovine plasma, and as such is more representative of the phenotypic effects of the venom as a whole, in comparison to the SVMP, SVSP, and PLA_2_ assays which are specific to one toxin family. All venoms demonstrated an overall procoagulant activity, as indicated by each of the curves initiating clotting earlier (within 4 minutes) and plateauing earlier (by 17 minutes) than the no-venom clotting control (initiated after 7 minutes and plateauing at ∼21 minutes) (Figure 3A). As with the other assays, there was variance between each venom with regards to the potency of the observed procoagulant effect; *B. moojeni*, *B. neuwiedi* and *B. atrox* were the most procoagulant and had caused complete clotting of the plasma by the second or third read timepoint (< 7 minutes, latter two venoms overlapped in kinetic reads). The venoms of *B. asper*, *B. jararaca* and *B. lanceolatus* exhibited comparable procoagulant profiles, demonstrating complete clotting by the fourth timepoint (∼ 10 minutes), while *B. alternatus* venom demonstrated the least potent procoagulant profile, with clotting plateauing around timepoint 6 (∼ 17 minutes), which was only marginally earlier than the no-venom control.

**Figure 3.**
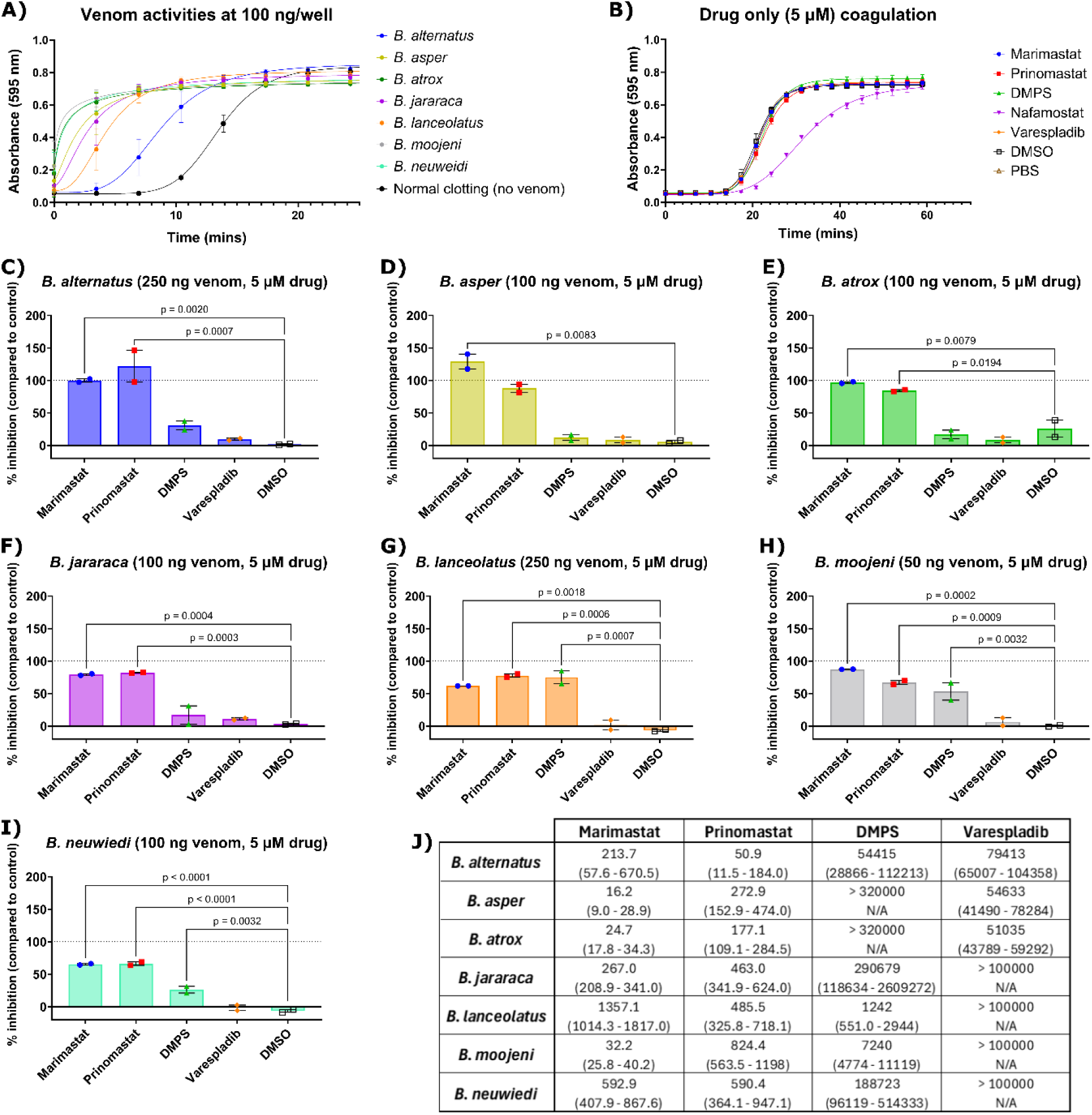
*In vitro* coagulation profile of seven different *Bothrops* species and inhibition of procoagulant activity by small molecule drugs. A) The coagulopathic profile of the seven *Bothrops* species at a comparable 100 ng dose in bovine plasma over 25 minutes at an absorbance of 595 nm at 25 °C (n=2, ± range). B) Coagulation profile in the absence of venom for SVMP inhibitors (DMPS, a metal chelator, and the MMP inhibitors marimastat and prinomastat), the PLA_2_ inhibitor varespladib and the serine protease inhibitor nafamostat. The profiles indicate no direct coagulopathy for any of the inhibitors except for nafamostat which has a strong anticoagulant profile at this dose (n=2, ± range). C-I) The percentage inhibition in the coagulation assay of the small molecules (excluding nafamostat due to the inherent anticoagulant activity) in each of the seven *Bothrops* venoms, tested at 5 µM (n=2, ± range) with a 25-minute preincubation at 37 °C. The adjusted dose for each venom indicated in the graph title was selected to provide comparable profiles for all seven venoms. Lines and p values indicate the significant differences determined by one-way ANOVA with Dunnett’s multiple comparisons test to the DMSO control (threshold p = <0.05). J) Dose response testing was performed for the four inhibitors in the coagulation assays and EC_50_s (nM) calculated using Prism v11 software (GraphPad) as presented this table (n=2, ±95% CI).

### *In vitro* drug inhibition

#### Inhibition of SVMP activity by MMPi and metal chelators

Following on from the SVMP assay in which variable activities were demonstrated across all venoms, a panel of 9 known matrix metalloproteinase (MMP) inhibitors were screened as dose response curves against a fixed dose of each venom with subsequent EC_50_ calculations to allow for potency comparisons (Figure 4A). Marimastat, a matrix metalloproteinase inhibitor (MMPi) that has demonstrated single digit nanomolar EC_50_s against a broad range of different snake venoms ^16,18,36,37^, was included as a gold standard. The top dose of marimastat, 10 µM, inhibited the SVMP activity of all venoms down to baseline (Figure 4B, p <0.0001) and demonstrated low nanomolar inhibition in the SVMP assay against most of the venoms tested (EC_50_ range 1.8 – 10.6 nM). DMPS, a heavy metal chelator that has recently completed a Phase I safety trial for snakebite indication ^22^, demonstrated sub-micromolar EC_50_s against all but two of the venoms (*B. asper =* 2019.0 nM, *B. atrox* = 2753.0 nM, with EC_50_s for the remainder ranging from 196.1 – 579.6 nM).

**Figure 4.**
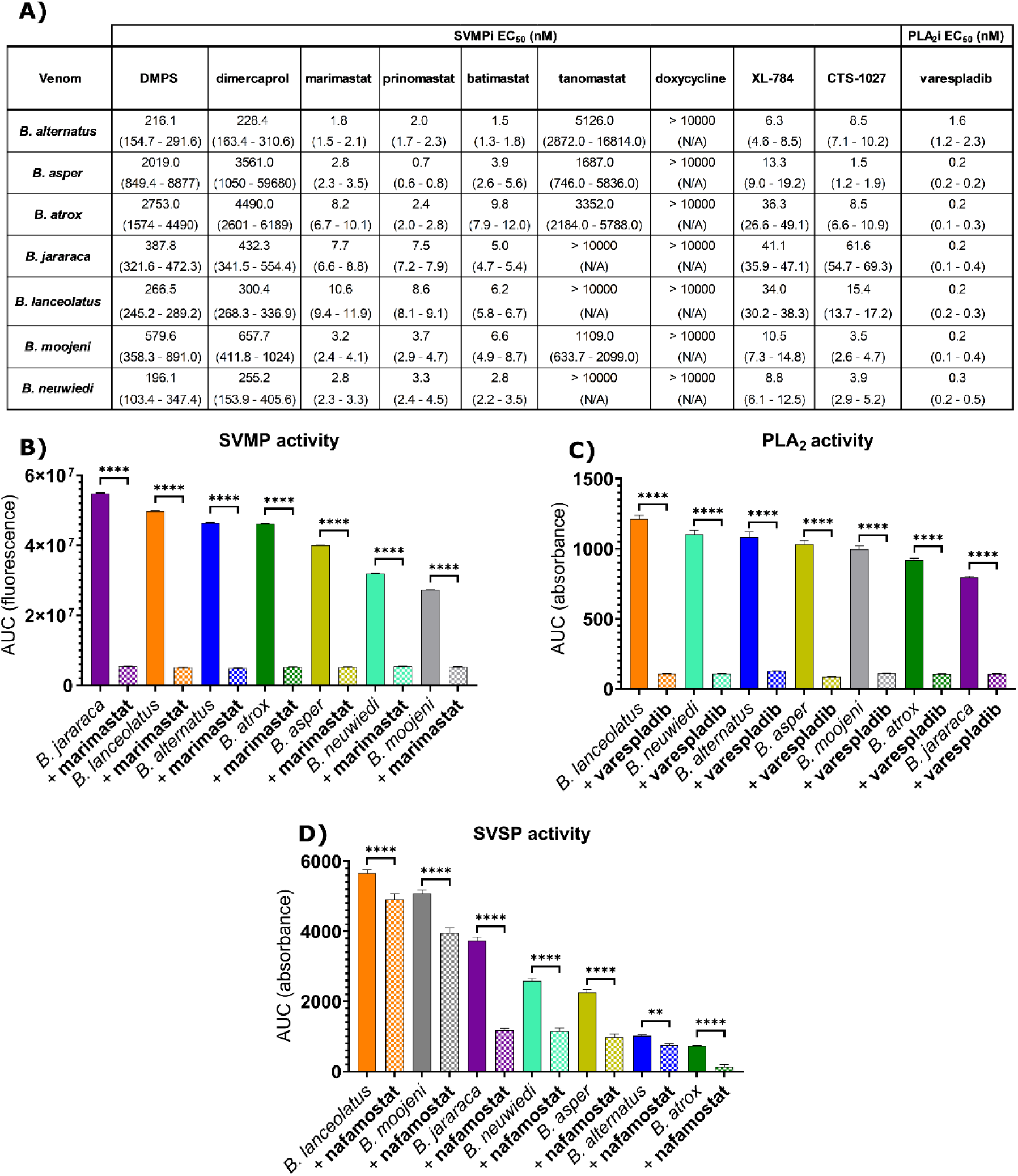
*In vitro* inhibition of seven different *Bothrops* species by small molecule drugs. The enzymatic activity of the seven *Bothrops* species with relevant inhibitors in the three toxin specific assays are presented, with the results analysed using area under the curve (AUC) and plotted using Prism version 11 software (GraphPad), with all drugs preincubated with the relevant drug for 25 minutes at 37° C. A) Dose response testing was performed in the SVMP and PLA_2_ assays using varespladib for the latter, but a wider panel of MMPis and metal chelators with prior evidence of SVMP inhibition in the SVMP assay. This allowed for the calculation of EC_50_s (with 95% CI shown) using Prism v10 software (GraphPad) which are displayed in this table (n=2, ± 95% CI). B) Snake venom metalloproteinase (SVMP) activity (1 µg venom per reaction) and inhibition by the MMP inhibitor, marimastat at 10 µM (AUC over 178 minutes, n=4, ±SE). C) phospholipase A_2_ (PLA_2_) activity (100 – 250 ng venom per reaction, depending on venom) and inhibition by the PLA_2_ inhibitor, varespladib at 10 µM (AUC over 13 minutes, n= ≥3, ±SE). D) snake venom serine protease (SVSP) activity (1 µg venom per reaction) and inhibition by the SP inhibitor, nafamostat at 10 µM (AUC over 35 minutes, n=4, ±SE). Statistical analysis of the data presented in panels B-D was assessed via two-way ANOVA with Šídák’s multiple comparisons test of each venom control compared to the matched venom treated with inhibitor (GraphPad Prism v11.0); ** = p = 0.0012, **** = p <0.0001.

These findings are in line with previously published data demonstrating the lower *in vitro* potency of DMPS when compared to MMPis due to their different mechanisms of action ^37,38^. Dimercaprol, another heavy metal chelator, demonstrated a pattern of inhibition in line with DMPS, albeit at slightly lower potency (EC_50_ range 228.4 – 4490 nM). The remaining six MMPis tested demonstrated comparable activity to previous studies ^38–40^. Prinomastat, batimastat, XL-784 and CTS-1027 demonstrated low nanomolar pan-species inhibition (EC_50_ = 1.5 – 15.4 nM across all four compounds) similar to marimastat, with the exception of CTS-1027 against *B. jararaca* (61.6 nM), and XL-784 against *B. lanceolatus*, *B. atrox,* and *B. jararaca* (EC_50_s of 34.0 nM for *B. lanceolatus*, 36.3 nM for *B. atrox*, and 41.1 nM for *B. jararaca*). *B. jararaca* and *B. lanceolatus* were not inhibited at any dose of tanomastat (EC_50_ > 10 µM), with the remaining venoms being inhibited at single digit micromolar (EC_50_ range = 1.1 – 5.1 µM). Doxycycline, a tetracycline antibiotic that has been shown to also target matrix metalloproteinases ^41^, and specifically SVMPs ^42^, was included as an unrelated compound family ^38^, though it displayed minimal inhibitory capacity against any of the venoms tested (all EC_50_s > 10 µM).

#### Inhibition of PLA_2_ activity by varespladib

Varespladib has previously been shown to potently inhibit PLA_2_ activity in a wide variety of venoms, with quoted EC_50_s in the sub-nanomolar range ^21,43^, as well as having been employed in a Phase II trial to determine efficacy against snakebite when given in combination with antivenom ^44^. In this study, varespladib mirrored the results seen in the literature, with pan-species inhibition at the top dose of 10 µM completely reducing PLA_2_ activity to baseline in all venoms (p <0.0001), and dose response EC_50_s in the sub-to single-digit nanomolar range (0.2 – 1.6 nM) (Figure 4A and C). Against most venoms, varespladib displayed similar EC_50_s of between 0.2 and 0.3 nM, while, perhaps surprisingly, *B. alternatus* demonstrated the highest EC_50_ of 1.6 nM despite having the lowest PLA_2_ activity of the tested venoms (Figure 2C). However, in an attempt to standardise the relative activities of the different venoms, we conducted these inhibition experiments using variable venom concentrations, so the presence of a higher concentration of venom (250 ng for *B. alternatus* compared with 25-50 ng for the other venoms) seems likely to explain the modest increase in EC_50_ observed for this species.

#### Inhibition of SVSP activity by nafamostat

The SVSP activity across the seven venoms was highly variable (Figure 2) and to a greater extent than the SVMP and PLA_2_ activities. Nafamostat is a serine protease inhibitor which has previously been shown to inhibit the SVSP activities of certain snake venoms at high doses (10 µM fully inhibited 1 µg venom activities ^37^). However, in this study for all seven *Bothrops* venoms at the same 1 µg of venom and 10 µM dose of nafamostat, although significant (Figure 4D, p < 0.0012) the inhibition was less than 80% of the SVSP activity.

This weak inhibitory activity justified no further assessment of EC_50_ testing, particularly given that higher dose testing at 80 µM resulted in only three venoms (*B. asper*, *B. atrox* and *B. jararaca*) being inhibited at greater than 80%.

#### Inhibition of coagulopathic venom toxins in bovine plasma

Based on the findings from the *in vitro* assays above, three representative SVMP inhibitors (marimastat, prinomastat and DMPS), the PLA_2_ inhibitor varespladib, and the SVSP inhibitor nafamostat were tested in the bovine plasma coagulation assay to determine whether any drug could restore normal coagulation in the presence of the various venoms. First, we assessed whether any of the compounds affected coagulation in the absence of venom (Figure 3B). While most had no effect, the serine protease inhibitor nafamostat was shown to be inherently anticoagulant at 5 µM and higher (Figure 3B) and so was excluded from downstream dose-response inhibition experiments. For the dose-response experiments, venom doses were adjusted to provide similar coagulation profiles for each venom to normalise the window between venom-induced clotting and no-venom clotting across all venoms (50-250 ng used). Marimastat and prinomastat displayed moderate to potent inhibition of all venoms at 5 µM (Figure 3C–3I, marimastat 61.8 to 129.1%, p = <0.0001 to 0.0083 and prinomastat 67.0 to 121.9%, p = <0.0001 to 0.0510), with a fairly broad EC_50_ range seen in both compounds (marimastat 16.2 to 1357.1 nM, prinomastat 50.9 to 824.4 nM; Table 3J). Though marimastat exhibited EC_50_s below 100 nM against more of the venoms (three vs one with prinomastat), it also demonstrated a high EC_50_ of 1357 nM against *B. lanceolatus*, a venom that prinomastat inhibited at a comparably lower EC_50_ of 485.5 nM. DMPS required a substantially higher dose for EC_50_ inhibition than any of the MMPis, which correlates with the lower potency of this compound in the SVMP assay. At a matched dose of 5 µM, DMPS was only weakly inhibitory against *B. alternatus, B. asper, B. atrox, B. jararaca* and *B. neuwiedi* venoms (Figure 3C-F: 30.7%, 11.9%, 16.8%, 17.0%, and 26.1% inhibition, respectively, with significant inhibition only observed with *B. neuwiedi*, p = 0.0032). Perhaps surprisingly, for *B. lanceolatus*, a venom that required a higher venom dose to match the procoagulant profiles of the other venoms (250 ng vs 50-100 ng), 5 µM of DMPS inhibited the venom by 75% (Figure 3G, p = 0.0007). In *B. moojeni*, 5 µM of DMPS inhibited the venom by 53% (Figure 3H, p = 0.0032). Interestingly, despite all venoms displaying procoagulant profiles, which could be assumed to be primarily driven by SVMPs, the PLA_2_ inhibitor varespladib demonstrated moderate inhibitory capacity of the procoagulant venom profiles at higher doses and was capable of generating EC_50_s for three of the venoms (*B. alternatus, B. asper,* and *B. atrox,* EC_50_s all > 50 µM, Figure 3J). However, at the matched 5 µM doses used for comparative purposes, varespladib had no effect on venom coagulopathy, suggesting this effect is predominately driven by SVMP toxins (Figure 3, panels C-I).

As marimastat displayed mediocre potency against *B. jararaca* venom in single-inhibitor experiments (79.2% inhibition at 5 µM and an EC_50_ of 267.0 nM in the coagulation assay, Figure 3), we postulated that the remaining procoagulant effect of the venom following inhibition with 5 µM marimastat could have been due to SVSP activity, as described for other species ^45,46^. To explore this, we used nafamostat in combination with marimastat, and also included *B. atrox* venom for comparison (96.7 % inhibition at 5 µM and an EC_50_ of 24.7 nM), alongside drug only controls to further assess the inherently anticoagulant effect of nafamostat at lower drug doses. As expected, the procoagulant effect of *B. jararaca* venom was not fully inhibited by 5 µM marimastat (64.9% inhibition), while 5 µM nafamostat only modestly inhibited procoagulant effects (13.0% inhibition) (Figure 5A and B). When combining the two inhibitors, an increase in venom inhibition was observed, with 5 µM doses of both drugs resulting in a percentage inhibition greater than the effect of marimastat alone (82.5% vs 64.9%, respectively with a significant difference between marimastat alone vs the combination of marimastat and nafamostat, p = 0.0009). This increase in inhibition is roughly equivalent to an additive effect of both compounds alone. Experiments with *B. atrox* venom showed that the procoagulant activity of this venom was near fully inhibited by 5 µM marimastat alone (96.4%), with similarly high inhibition at 2.5 µM (89.4%) and 1.25 µM (83.5%) (Figure 5C and D), whilst nafamostat displayed no effect on the procoagulant venom profile, even at the 5 µM drug dose shown to be anticoagulant (153.9% drug only inhibition, Figure 5E and F), indicating that the anticoagulant effect of this compound cannot alter the procoagulant effects of the SVMPs present in the venom. When combining 5 µM marimastat and nafamostat with *B. atrox* venom, the resulting coagulation profile shifted to being anticoagulant (i.e. >100% inhibition, 136.6%, with a significant difference between marimastat alone vs the combination of marimastat and nafamostat, p = < 0.0001), likely the result of the anticoagulant effects of nafamostat being realised once procoagulant SVMPs were inhibited by marimastat.

**Figure 5.**
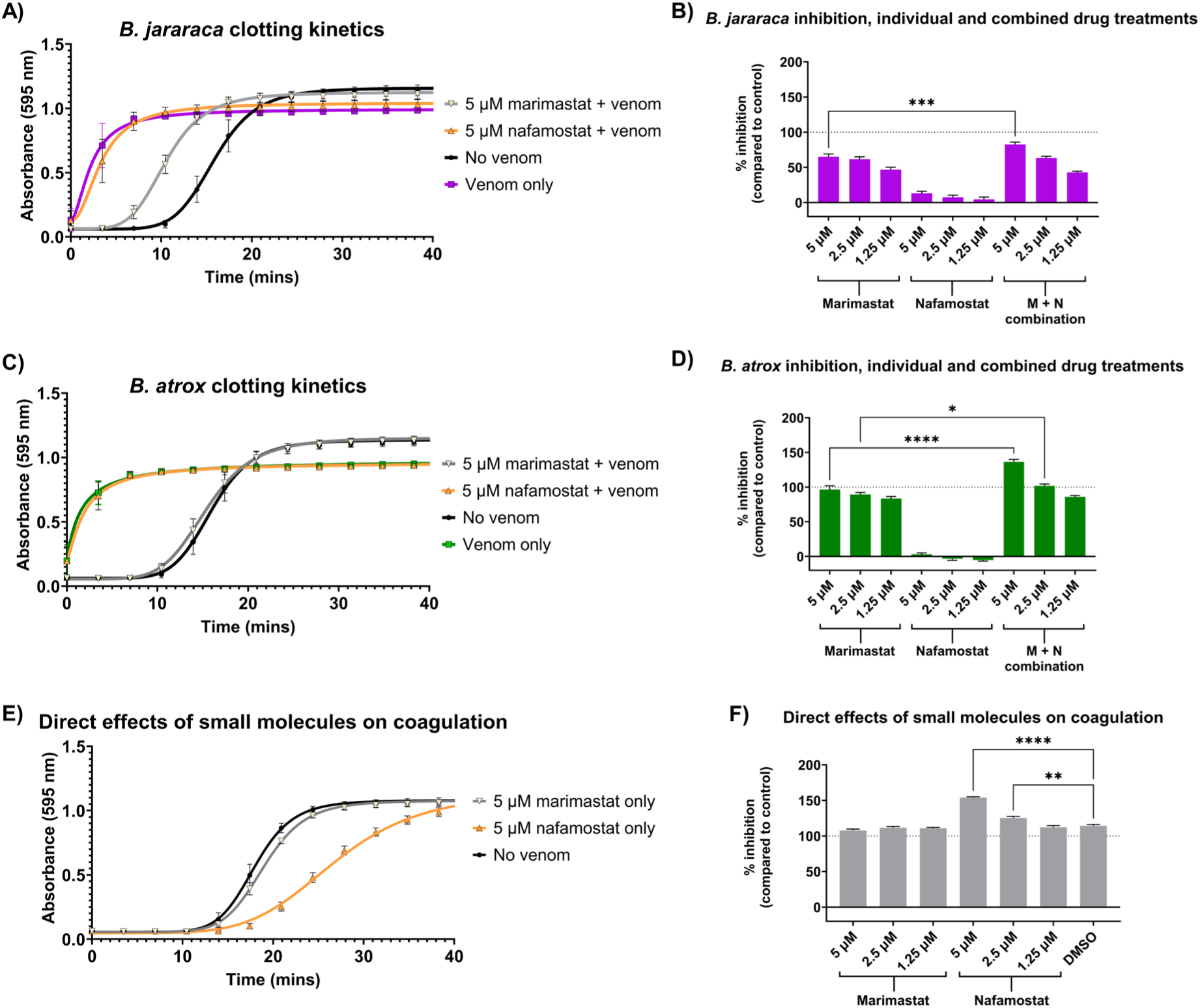
*In vitro* coagulation profile of two coagulopathic *Bothrops* species and inhibition by combination therapy of two small molecule drugs. The effect of *Bothrops jararaca* (A-B, purple) and *Bothrops atrox* (C-D, green) with and without preincubated small molecule inhibitors is presented alongside the direct effect of these inhibitors (E and F) in the coagulation assay. (A and C) The kinetic curves demonstrate the procoagulant effect of both venoms, compared to normal clotting control (black circles). The SVSP inhibitor nafamostat (orange triangles) fails to rescue the procoagulant effect, whilst marimastat (5 µM, white triangles) fully inhibits the procoagulant effect of *B. atrox* and partial rescue of *B. jararaca* (n=6, ± SD). (B and D) Percentage inhibition of coagulopathy by *Bothrops* species in single treatment by small molecule inhibitors (n=6, ± SE) matched that of the kinetic curves for the top 5 µM dose of marimastat with subsequent lower doses slightly reducing the percentage inhibition, nafamostat resulted in minor inhibition at any doses. (E and F) Combination testing of marimastat and nafamostat, for both venoms appears to reflect an additive effect, however in the no-venom control (E, kinetic curve; F, % inhibition; n=12, ± SE, nafamostat shows a dose-dependent increase beyond 100% in the absence of venom, demonstrating that it is directly anticoagulant at higher doses. Marimastat has no effect on plasma clotting, indicating that the effects seen in graphs (A to D) are through inhibition of venom. Statistical analysis of the data presented in panels B,D and F was assessed via one-way ANOVA with Šídák’s multiple comparisons test, * p = 0.0163, ** p = 0.0047, *** p = 0.0009, **** p = <0.0001.

### Inhibition of coagulopathic venom toxins by human whole blood thromboelastography

To further investigate the plasma coagulopathy results in a more clinically relevant system we spiked *B. atrox* and *B. jararaca* venom into human whole blood and evaluated the resulting coagulation profiles using thromboelastography ^26^. The methodology utilised in this study involved the addition of venom, with or without marimastat, to calcium chloride and whole blood immediately before reading on a ROTEM instrument. Pilot experiments with nafamostat only revealed direct anticoagulant effects (clotting time [CT]: 3600 at 5 µM and 1368 seconds at 1.25 µM, compared to 682.7 for 5 µM marimastat only and 644.3 seconds for vehicle control), and thus was excluded from further study. As in the bovine plasma assay, both *B. atrox* and *B. jararaca* venom presented with a rapid clotting time (mean CT: *B. atrox,* 133.3s; *B. jararaca*, 184.3s; vs no venom control, 664.3s, both p <0.0001 compared to no venom control) and similar clotting strength (mean maximum clot firmness [MCF]: *B. atrox,* 68.3; *B. jararaca,* 66.7; vs no venom control, 58.0, p = 0.0038 and 0.0149, respectively, compared to no venom control) (Figure 6). The reduced CT time caused by both venoms was partially inhibited by marimastat, resulting in restoration of 67% and 62% of the longer CT of the no venom control against *B. atrox* and *B. jararaca* venom, respectively (both p = <0.0001). Despite this similar inhibition of the CT, there was a notable difference in the ability of marimastat to reduce the increased clotting strength induced by the two venoms, with *B. jararaca* venom inhibited by 74% compared with 29% with *B. atrox* though neither reduction was statistically significant (p > 0.068). Due to the previously described interfering effect of nafamostat in this assay, we were unable to explore whether parallel inhibition of SVSPs might further restore the clotting strength induced by *B. atrox* venom closer to baseline, though data from the previously described bovine plasma assay (Figure 5) suggested little effect by nafamostat for this venom, hinting that perhaps other toxins families might be responsible for this phenomenon.

**Figure 6.**
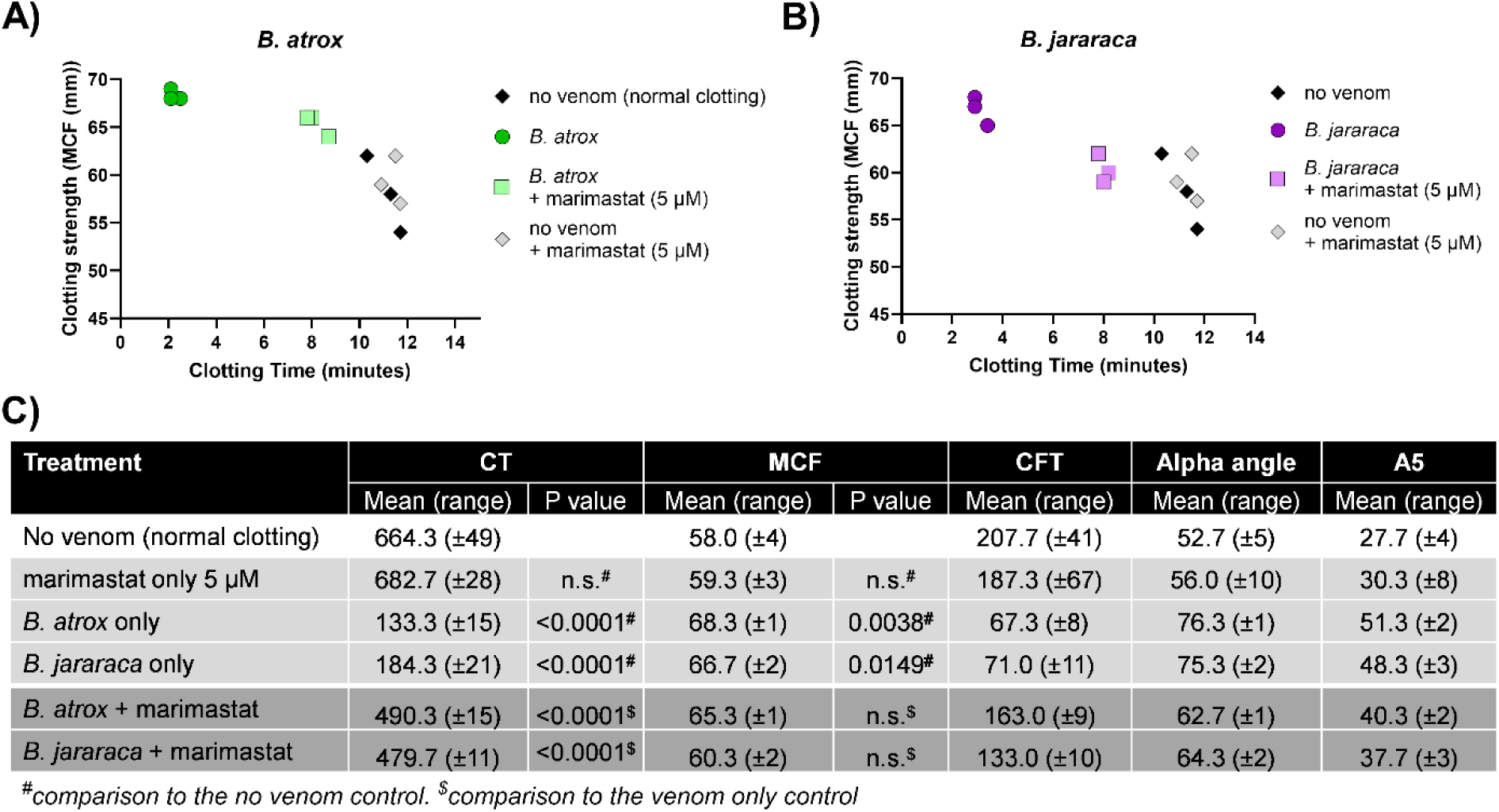
Thromboelastography (TEG) profiles of two representative *Bothrops* species and inhibition by the SVMP inhibitor marimastat. A and B) Thromboelastography profile of respectively *B. atrox* and *B. jararaca* using an XY plot of clotting time vs clotting strength. Both venoms (0.6 µg per 300 µL reaction) have strong procoagulant activity (circles) compared to the no venom controls (black diamond) or drug only control (grey diamond). Marimastat inhibits the procoagulant activity of both venoms (squares) (n >3). C) The parameters reported by the ROTEM instrument are displayed including the clotting time (CT; seconds), maximum clot firmness (MCF; mm), clot formation time (CFT; seconds), alpha angle (°) and amplitude at 5 minutes (A5; mm), each is the mean of the triplicate results ± the related range. Independent statistical analysis of CT and MCF were performed against either the no venom^#^ (pale grey rows) or venom only controls^$^ (dark grey rows) using one-way ANOVA with Tukey’s multiple comparisons test. n.s. = not statistically significant.

### Inhibition of venom lethality in an *in vivo* chicken egg model of envenoming

To further evaluate the potential protective effects of marimastat seen in the coagulation assay above, we investigated *in vivo* protection in an insensate chicken egg model of snakebite envenoming. Using *B. atrox* venom as a model, we first demonstrated that topical application of venom to the vitelline vein of embryos at a dose of 20 µg resulted in clear observable pathology, characterised by rapid destruction of the vasculature within 1 hour (Supplementary Figure 1), and lethality by the end of the experimental time course of 6 hours (80% lethality, n=20). Topical treatment with marimastat (0.5 µg, 1.0 µg and 5.0 µg, n=5 per group) immediately after venom dosing resulted in dose-dependent increases in efficacy against vasculature damage and venom-induced lethality, with the highest dose providing complete protection against lethal venom effects (0.5 µg, 60% survival, p = 0.0498; 1.0 µg, 80% survival, p = 0.0306; 5.0 µg, 100% survival, p = 0.0130) (Figure 7 and Supplementary Figure 1). To contextualise these findings gained with the MMPi marimastat, we repeated these experiments using the lead SVMP-inhibiting metal chelator, DMPS, at the same therapeutic doses. The lowest dose of DMPS (0.5 µg) resulted in observable venom pathology and a survival curve near identical to the venom control (20% survival), indicative of no protection (Figure 7 and Supplementary Figure 1). However, higher doses of DMPS (1.0 and 5.0 µg) showed clear evidence of venom neutralisation and provided complete protection against venom-induced lethality (100% survival, both p = 0.0130) (Figure 7 and Supplementary Figure 1). These findings further emphasise the promising preclinical potency of this drug ^37^, despite its reduced *in vitro* SVMP inhibitory potency in comparison with MMPis.

**Figure 7.**
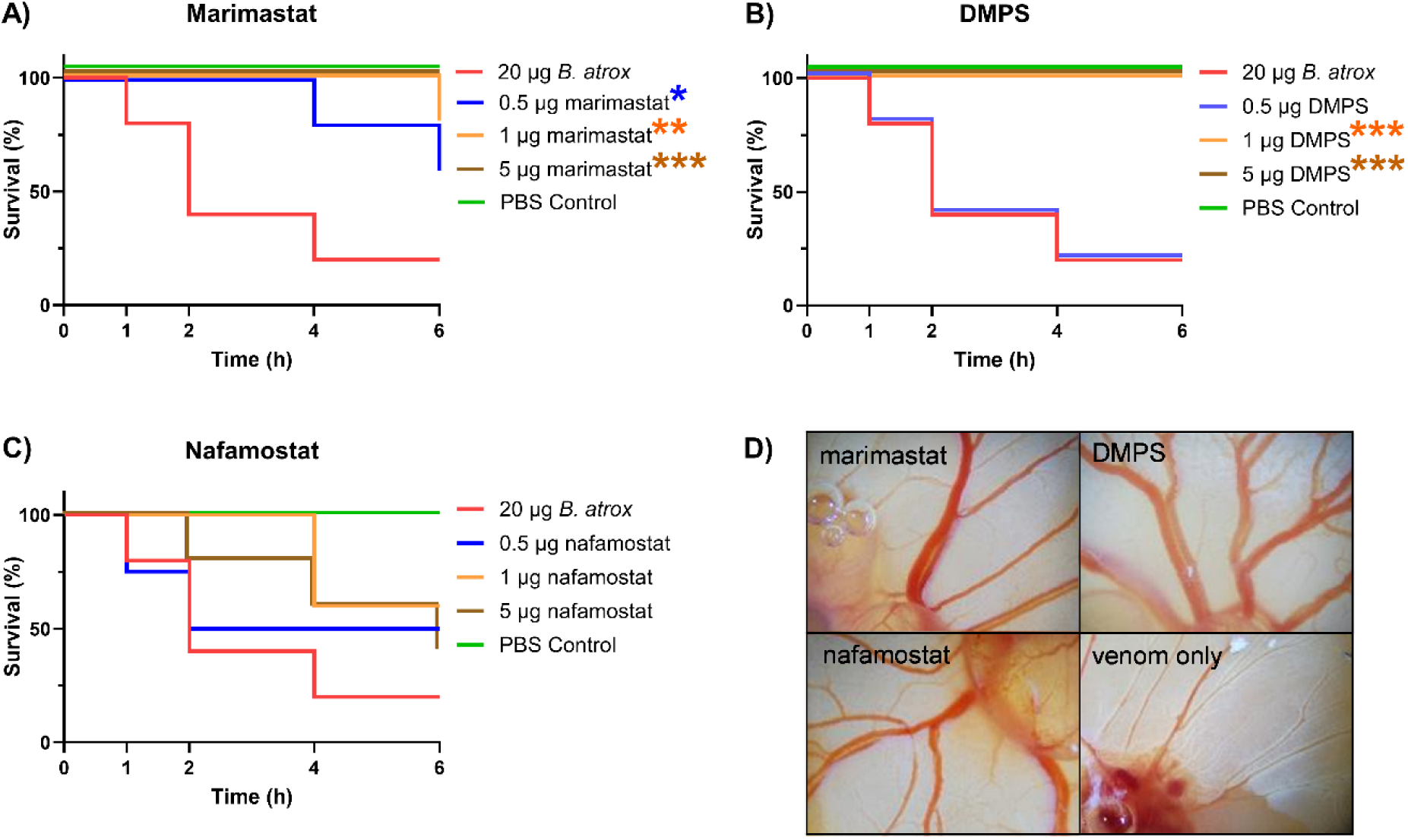
Survival curves of chicken embryos dosed with *B. atrox* venom with and without representative small molecule drugs. Groups of 6-day post-fertilisation chicken egg embryos were dosed on to the vitelline vein with either PBS (negative control) or *B. atrox* venom (20 µg) with or without a subsequent dose of 0.5 µg, 1 µg and 5 µg of A) marimastat, B) DMPS and C) nafamostat. All treatment group sizes were n=5, the PBS control contained an n=12, while the venom only control contained an n=20. D) Representative pathology images of chicken embryos 1 hour post-dosing with *B. atrox* venom with and without 5 µg of the inhibitory drugs, showing varying degrees of protection against the vascular damage shown in the venom only control (see also Supplementary Figure 1). Log-rank test (Mantel-Cox) with Holm-Šídák’s multiple comparisons test was carried out for all treatments against the relevant venom-only control and significant differences are indicted by asterisks found to the right hand side of the relevant legends in each panel; * p = 0.0498, ** p = 0.0306 and *** p = 0.0130.

Finally, to further investigate whether inhibition of SVSP toxins might also be beneficial for *in vivo* protection against *Bothrops* envenoming, we repeated these *B. atrox* chicken embryo experiments using the SVSP inhibitor nafamostat. As outlined above, due to its potent off target anticoagulant effects, we were unable to robustly evaluate the value of inhibiting SVSP toxins with nafamostat in the coagulation assays, but control doses of nafamostat alone (5.0 µg) had no observable effect on the embryos (100% survival, Supplementary Figure 1), facilitating use in these haemotoxicity and lethality focused experiments. Perhaps unsurprisingly, embryos dosed with venom and nafamostat showed evidence of pathology at all tested drug doses (0.5, 1.0, 5.0 µg), although at the end of experiment six hours post-dosing, some evidence of modest efficacy was observed. Dosing with nafamostat resulted in 40-60% embryo survival across the three dose groups, which compared favourably with the 20% survival reported in the venom only control group. However, statistical analysis demonstrated these differences were not significant (p = 0.139-0.414). These findings suggest that for these venoms inhibition of SVSPs is likely of secondary importance when compared with SVMP inhibition, which can convey complete protection from venom-induced lethality.

## Discussion

Using a wide range of *in vitro* approaches, including venom composition profiling by SDS-PAGE and toxin specific activity assays, we show that *Bothrops* species have highly variable SVMP, SVSP and PLA_2_ activities (Figure 2 and 3), agreeing with the widely reported literature describing their variable interspecies venom compositions (Figure 1). For example, *B. alternatus, B. asper* and *B. atrox* had low SVSP activities, comparably high SVMP activities, yet highly variable PLA_2_ activities. Conversely, *B. moojeni* and *B. neuwiedi* presented with the lower range for SVSP activity, but had variable SVMP and PLA_2_ activity. Crude comparisons between the literature reports of proteomic venom composition (Figure 1) to the enzymatic activity of the venoms in this study show similar rankings for *B. alternatus* and *B. atrox* (*B. alternatus*: low PLA_2_, low SVSP and high SVMP. *B. atrox*: variable PLA_2_, low SVSP and high SVMP), but there was limited correlation for the remaining venoms. This lack of correlation is not surprising as the literature clearly describes evidence of variation in venom composition within individual *Bothrops* species caused by various factors, such as venom sampling location inclusive of habitat variation, gender or ontogeny differences and wild caught versus captive bred (Supplementary Table 1) ^5,7,8,47,48^. It should further be noted that the data we show for each species in Figure 1 are not uniform, with some species represented by the venom composition based on minimal publications sampling a small pool of individuals, while other species have multiple publications comprising analysis of multiple pools of venom from diverse individuals (Supplementary Table 1). Future proteomic characterisations of the specific venom samples used in this study, which were all sourced from a historical collection (except for *B. lanceolatus*), would be informative in this regard.

Despite the variation in toxin activity across the *Bothrops* species selected in this study, we observed pan-species nanomolar neutralisation at the enzymatic level for two small molecule drugs entering snakebite clinical trials, and which neutralise SVMP (marimastat) and PLA_2_ (varespladib) toxins (Figure 4). The second lead SVMP inhibiting drug, the metal chelator DMPS, has reduced *in vitro* neutralising potency across the venoms tested (EC_50_ ∼200 nM – 3 µM). However, the serine proteinase inhibitor nafamostat required greater than 10 µM to achieve full neutralisation against any of the venoms. SVSP toxins are present in the venoms of various *Bothrops* species, can be of comparable abundance to SVMP toxins in certain species ^7,49^, and had high enzymatic activity in the *B. jararaca*, *B. lanceolatus*, *B. moojeni* and *B. neuwiedi* venoms tested here (Figure 2). Despite the potentially important coagulopathic effect of SVSPs on the haemostatic system, small molecule inhibitors of SVSPs have received limited attention, in contrast to the recent drug discovery efforts activity against PLA_2_ and SVMP toxins. Here we used the previously described, repurposed, serine protease inhibitor nafamostat, which was previously used in small molecule combination therapies to investigate its value in preventing venom lethality preclinically ^37^.

However, nafamostat showed limited additive value compared to a combination of SVMP and PLA_2_ inhibitors in that study, while the low *in vitro* inhibitory potency of the drug described here (>10 µM EC_50_) coupled with its innate anticoagulant nature via interactions with cognate serine proteases such as thrombin ^50^, severely hampered our assessment of the role SVSPs play in the coagulation disturbances caused by *Bothrops* venoms in the plasma coagulation and whole blood TEG assays. Despite the lack of evidence we observed, previous studies suggests that SVSPs are likely important pathological components of *Bothrops* venoms^51^. There therefore remains a strong rationale for future drug discovery activities to identify novel broad-spectrum inhibitors against SVSP toxins, with a focus on optimising their potency and selectively to avoid off target effects against mammalian serine proteases involved in the coagulation cascade ^52^.

Viperid PLA₂ toxins induce coagulopathy primarily through their anticoagulant activity. Although we identified clear enzymatic activity of PLA_2_ across many of our sampled species, we observed no anticoagulant activity in either the plasma or whole blood experiments, even when procoagulant SVMP toxins were inhibited. Despite this lack of coagulopathic effect, the potent pan-species inhibitory activity of varespladib observed in the PLA_2_ enzymatic assay (EC_50_s ranging from 0.14 to 1.45 nM) is likely still to be beneficial at inhibiting non-coagulant PLA_2_-associated pathology, such as myonecrosis, tissue damage and inflammation ^43,53^.

Indeed, myonecrosis, caused by myotoxic PLA_2_s, is a pathology commonly associated with *Bothrops* envenoming ^54^. Both enzymatically active and inactive myotoxins have been described in several *Bothrops* venoms and varespladib has been previously shown to neutralise their activity ^54–56^. Our data further support the broadly neutralising activity of varespladib against enzymatically active PLA_2_, which may include myotoxins, however a limitation of our study was that we did not conduct assays to specifically evaluate the neutralising effects against myotoxicity. The potential value of a PLA_2_ inhibitor against *Bothrops* venoms is further highlighted by prior reports of poor antivenom performance against certain PLA_2_ isoforms due to either their lack of inclusion in the immunisation mixture or due to poor tissue penetration of the antivenom ^25,31,35^.

Challenges associated with the neutralising efficacy of antivenoms, such as Instituto Butantan’s polyvalent antivenom antibotrópico, have also been reported against certain SVMP isoforms ^31,35^. The pan-species activity of MMPis like marimastat is widely reported to be facilitated by broad spectrum inhibitory effects against varying isoforms of SVMPs ^38,57^. In our studies, marimastat demonstrated pan-species inhibitory activity across diverse venoms with nanomolar EC_50_s in the SVMP enzymatic assay against all venoms (1.8 to 10.6 nM) and in the plasma coagulation assay against all but one venom (16.2 to 592.9 nM, except for *B. lanceolatus*, 1357.0 nM). Although both marimastat and prinomastat exhibited similar potency in both *in vitro* assays (prinomastat EC_50_ range 0.7-8.6 nM in SVMP assay, 50.9 - 824.4 nM in coagulation assay) and both these repurposed drugs have undergone clinical evaluation for other indications, marimastat has been more extensively characterised preclinically than prinomastat for snakebite, and will soon enter a Phase II clinical trial evaluation for this indication ^58^. Marimastat also has a longer half-life in humans of 8-10 hours ^59^, compared to prinomastat (2-5h ^60^) further justifying the prioritised progression. The *in vitro* results were further supported via the use of TEG assessment of coagulopathy in human blood, where the distinct venoms of two WHO category one (highest medical importance) species, *B. atrox* and *B. jararaca*, ^61^. In Brazil, *B. atrox* is implicated in 80-90% of snakebites^24^, but this species also extends into the Amazonian regions of many other countries (Figure 1A), while *B. jararaca* is found in the southern states of Brazil, northeastern Paraguay and northern Argentina, where it is a leading cause of snakebite, especially in the densely populated areas of southeastern Brazil^7^. Despite their differing *in vitro* profiles, in the TEG human whole blood assay, marimastat rescued the procoagulant activity of *B. atrox* and *B. jararaca* venom by inhibiting the induced rapid clotting time and increased clotting strength, reverting the clotting profile towards normal parameters seen in the absence of venom. Further, marimastat provided protection in an *in vivo* model of haemotoxicity, with chicken egg embryos dosed with *B. atrox* venom fully protected against lethality and the vascular destructive effects caused by this venom at the 5 µg therapeutic dose tested (vs 20 µg venom). These findings provide further confidence in the potential therapeutic value of marimastat, which remains a lead candidate repurposed drug currently approaching clinical development. Future evaluation of its safety and efficacy in a Phase II clinical trial against *B. atrox* envenomings will be revealing.

As seen in previous work, there was clear superiority in terms of the *in vitro* SVMP inhibitory potency of matrix metalloproteinase inhibitors over metal chelators (Figure 4A) ^38^.

Marimastat and prinomastat exhibited EC_50_s in the low nanomolar range (from 0.7 nM for prinomastat vs *B. asper* to 10.6 nM for marimastat vs *B. lanceolatus*) in the SVMP assay, resulting in at least a 20-fold increase in potency, and often >100-fold potency, over the lead metal chelator DMPS (EC_50_ range of 196.1 to 2019 nM). A similar trend was observed in the coagulation assay (Figure 3J; EC_50_ range of 16.2 to 1357.0 nM for marimastat and 50.9 to 824.4 nM for prinomastat vs 1242 nM to >320,000 nM for DMPS), although the anti-coagulopathic potency of DMPS was dramatically higher against *B. lanceolatus* venom than all other venoms tested (1242 nM) and highly comparable to the EC_50_ of marimastat against this venom (1357 nM). Given that envenomings by *B. lanceolatus* are known to often be somewhat distinct to bites by other *Bothrops* species, including presenting with severe thrombotic complications ^62^, further evaluation of the potential protective effects of the already licensed drug DMPS in a preclinical setting against this species would be of great interest. Finally, despite the considerable differences in *in vitro* potency mentioned above, DMPS provided highly comparable protective effects to marimastat against the *in vivo* pathology caused in chicken egg embryos by *B. atrox* venom. In this pilot study measuring *in vivo* haemotoxicity, treatment with 1 µg and 5 µg of marimastat resulted in 80% and 100% protection against lethal venom effects, while DMPS was fully protective at both these doses. These findings further highlight the apparent *in vitro*-*in vivo* potency disconnect previously described for this metal chelator ^63^ and highlight that caution should be applied when triaging SVMP inhibitors with different mechanisms of action to MMPis based solely on *in vitro* enzymatic inhibitory potency.

This manuscript reinforces prior work demonstrating that the diversity of venom compositional and functional potency found across the *Bothrops* genus of snakes is substantial. Here we evidence this variation in a comparative manner, highlighting both similarities and differences in the enzymatic activity (SVMP, SVSP and PLA_2_) and coagulopathic (*in vitro* bovine plasma coagulation and TEG human whole blood) effects of various *Bothrops* venoms. Despite such variation, we show that three of the leading small molecule treatments currently progressing into clinical trials (marimastat, DMPS and varespladib) can collectively neutralise all seven venoms in terms of their enzymatic PLA_2_ and SVMP activity, while the SVMP inhibitors were also effective against the action of coagulotoxins and provided *in vivo* protection against lethal haemotoxicity in a chicken embryo model. While the current study did not assess the efficacy of these molecules against *Bothrops* venoms in a murine preclinical model of envenoming, previous work suggests the chicken embryo model can be highly informative and translatable relative to the standard WHO mouse preclinical model ^64–66^, with correlations between the two previously noted ^67,68^. The findings presented here therefore strongly advocate for onward progression of the described protective drugs into murine models in the future, with the priority to evaluate whether *in vivo* protection remains in rescue models of oral drug dosing and, if successful, to determine an appropriate dosing regimen that confers said protection.

Additional future priorities include: i) evaluating the diversity of toxin isoforms that are inhibited by these specific drugs, including those described to have limited neutralisation by available antivenoms ^9,25,35^, ii) undertaking drug discovery activities to identify novel SVSP inhibitors specific to venom toxins, and iii) as mentioned above, progressing lead repurposed drugs and drug combinations into conventional murine preclinical models ^11^ to evaluate their efficacy against both the systemic and local signs of envenoming *in vivo*. Overall, this study provides convincing evidence of the potential value of small molecule-based toxin inhibitors for the treatment of snakebite in the Neotropics, adding further weight to the recent paradigm shift towards early therapeutic interventions via oral dosing in community settings ^15,22,44^.

Indeed, certain settings within the Neotropics provide exciting potential for the robust, future evaluation of the efficacy of small molecule drugs against snakebite - in particular the use of SVMP inhibitors against *B. atrox* - and the outcomes of future clinical trials have the potential to provide valuable proof of concept for the future translation of safe and effective oral snakebite drugs.

## Author contribution

R.H.C., N.R.C., A.W. and S.K.M. conceptualised the project and wrote the manuscript, with funding acquired by N.R.C. The *in vitro* bioassays were performed by A.W., R.H.C. and S.K.M. The *in vivo* embryo assays were performed by E.S., L-O.A. and T.D.K. All authors analysed the results and reviewed the manuscript.

## Acknowledgements

We gratefully acknowledge MicroPharm Limited for provision of *Bothrops lanceolatus* venom. We also thank Camille Abada and Iara Cardoso for venepuncture and anonymised donors for their provision of blood samples. Our thanks are extended to Dr. Charlotte Dawson for the establishment of the chicken egg model and training the authors on its use. This work was funded by Wellcome (#221712/Z/20/Z to N.R.C.).

## Methods

### Venoms

Representative venoms were selected to cover the diversity of the *Bothrops* genus. All, except for *Bothrops lanceolatus* (which was gifted by MicroPharm Limited), were historical samples sourced from the herpetarium facility at the Centre for Snakebite Research and Interventions (CSRI) at the Liverpool School of Tropical Medicine (LSTM). Due to the historic nature of the venom samples, the source locality is not available beyond country of origin, with the exception of *B. lanceolatus* which is endemic to Martinique. The species were: *Bothrops alternatus* (Brazil), *B. asper* (Costa Rica – Atlantic), *B. atrox* (Colombia), *B. jararaca* (Brazil), *B. lanceolatus* (Martinique), *B. moojeni* (Brazil) and *B. neuwiedi* (Brazil).

Crude venoms were stored lyophilised at 2–8 °C before reconstitution to 10 mg/mL in sterile Phosphate Buffered Saline (PBS, pH 7.4) (Gibco, Cat.no. 10010023) prior to use.

### Drugs

The small molecule drugs used in this study were selected based on their previously reported inhibitory activity against snake venom SVMP, PLA_2_ or SVSP toxins ^37,38,43,57^. The SVMP-inhibiting matrix metalloproteinase (MMP) inhibitors were sourced from MedChemExpress - prinomastat hydrochloride (Cat. no. HY-12170A), XL-784 (HY-19485, 98.25%) and CTS-1027 (HY-10398, 99.24%); Sigma-Aldrich - marimastat (Cat. no. M2699), batimastat (Cat. no. SML0041) and doxycycline (Cat. no. D9891); and Cayman chemicals - tanomastat (Cat. no. 9258). The SVMP inhibiting metal chelators were dimercaprol (Cat. no. 64046, Sigma-Alrich) and DMPS (Cat. no. H56578, Alfa Aesar). The PLA_2_ inhibitor was varespladib (Cat no: SML1100, Sigma) and the SVSP inhibitor was nafamostat mesylate (Cat. no. ab141432, Abcam). All drugs were resuspended in dimethyl sulfoxide (DMSO) (Cat. no. D2650-100ML, Sigma-Aldrich).

### SDS-PAGE gel electrophoresis

Five micrograms of each venom were mixed in an equal volume of 2 X sample loading buffer (62.5 mM Tris-Cl pH 6.8, 25% v/v glycerol, 2% SDS, 0.75% bromophenol blue) with 100 mM dithiothreitol, incubated at 100°C for 5 minutes, before loading on a 4–20% Mini-Protean TGX gel (BioRad, Cat. no. 456-1096) with the addition of a molecular weight marker (5 µL, PageRuler Prestained Protein Ladder, Thermo, Cat. no. 26616) to one lane of the gel. After electrophoresis at 200 V for 30 minutes, the gel was stained with Coomassie blue (50% methanol, 40% deionized water, 10% glacial acetic acid and 0.1% Coomassie Brilliant Blue) for 1 hour at room temperature with gentle shaking, then de-stained for 2 hours at room temperature in 50% methanol, 40% deionized water and 10% glacial acetic acid. Gels were rinsed in deionised water and imaged on a GelDoc (Bio-Rad) under white light.

### SVMP *in vitro* assay

The SVMP assay utilises a quenched fluorescent substrate for MMPs previously utilised to assess the activity of SVMPs ^36–38,57^. One microgram per well of venom was added to a flat-bottomed 384-well plate (Greiner, Cat. no. 781101) in 15 µl PBS, before incubation at 37°C for 25 minutes (to keep conditions identical to the drug inhibition assays described below). Plates were then allowed to acclimatise to room temperature for 5 minutes, before 75 µl of fluorescent substrate (Bio Techne, Cat. no. ES010) at a final 7.5 µM reaction concentration in SVMP assay buffer (50mM Tris HCl pH 7.5, 150mM NaCl) was added to each well.

Immediately following addition of substrate, the fluorescence (excitation 320 nm, emission 420 nm) was read kinetically on a CLARIOstar Plus (BMG labtech) at 25°C. Venom inhibition assays were conducted in the same manner, with an additional preliminary preincubation step in which various concentrations of the drugs (final dose range of 10 µM – 0.17 nM) were created in DMSO. These were then stamped as 0.91 µL droplets at 100x the desired final concentration (final well volume being 91 µL) onto flat-bottomed 384-well plate wells prior to venom addition, allowing drug-venom interaction prior to substrate addition. For both protocols the AUC over 108 minutes was calculated for every condition to quantify activity. This time frame was selected based on prior knowledge of active venoms having capacity to fully convert the substrate ^36–38,57^. For the inhibitor testing the AUC was converted to a percentage of venom inhibition by normalising to the negative and positive controls.

These values were then plotted as a dose-response curve, and EC_50_ values calculated. Statistical analysis of the inhibitors was assessed using two-way ANOVA with Šídák’s multiple comparisons test of each venom control compared to the matched venom treated with inhibitor.

### PLA_2_ *in vitro* assay

The PLA_2_ assay uses a commercially available colorimetric assay kit (Abcam, Cat. no. ab133089) and relies on the cleavage of dithiol groups from a substrate (PLA2 Diheptanoyl Thio-PC) by PLA_2_s. A third component, DTNB, is added to the reaction, and binds to the freely available thiol groups, producing a colorimetric change. This kit has been adapted for venom PLA_2_ activity assessment in a 384-well format ^37,43^. For venom activity assays, 100 ng of each venom in a 10 µL volume were added to appropriate wells of a flat-bottomed 384-well plate (Greiner, Cat. no. 781101) then incubated for 25 minutes at 37°C for consistency with later dose-response experiments. Following incubation, the plates were allowed to acclimatise to room temperature prior to addition of 5 µL DTNB (resuspended in H_2_O) and 30 µL of PLA_2_ substrate (diluted in supplied assay buffer), and the plates read kinetically using an absorbance protocol (405 nm) on a CLARIOstar Plus at 25 °C. Thereafter, for each venom, the highest venom dose that gave a linear increase in absorbance over the 15-minute read time was selected for drug inhibition studies, in which dose-response curves of varespladib (final concentration range of 11.1 µM – 1.11 pM) were created in DMSO and first stamped onto a flat-bottomed 384-well plate as 0.5 µL droplets at 90x the desired final concentration (45uL final well volume) to allow for drug-venom preincubation prior to substrate addition. The calculated AUCs over 12 minutes (based on the Abam guidelines) were normalised to a percentage inhibition (compared to the positive and negative controls) and plotted as dose response curves, allowing for EC_50_ to be calculated. Statistical analysis of the inhibitors was assessed using two-way ANOVA with Šídák’s multiple comparisons test of each venom control compared to the matched venom treated with inhibitor.

### SVSP *in vitro* assay

The serine protease activity of the venoms was tested using a commercial chromogenic broad-spectrum peptide substrate (S-2288, Quadratech Diagnostics Ltd) to quantify the cleavage of the substrate by serine protease via absorbance, as previously described ^37^. The substrate was diluted in water to make a 6 mM stock solution. The reaction was performed with the addition of venom, buffer and substrate at a volume ratio of 1:1:1 with a final volume of 45 µL per well of a 384-well plate (Greiner, Cat. no. 781101). For each of the venoms, 1 µg was added per well (15 µL of 0.07 µg/µL diluted in PBS from a 10 mg/mL stock) prior to the addition of 15 µL of buffer (100 mM Tris pH 8.5, 100 mM NaCl) and incubated for 30 minutes at 37 °C. Following this incubation, 15 µL of the substrate stock solution was added per well (2 mM final concentration) and absorbance immediately measured kinetically at 405 nm on a CLARIOstar Plus plate reader at 37 °C. Drug activity was investigated identically to above, with an additional preliminary preincubation step in which dose-response curves were created in DMSO, via a 12-point curve ranging from 80 µM to 39 nM in 1:2 dilution steps. From each dose 0.5 µL was added to the venom in each relevant well and preincubated for 30 minutes at 37 °C before the buffer and substrate was added. For analysis, the AUC was calculated over a 35-minute time interval and all data normalised by subtracting the no venom control (PBS). This time frame was selected based on prior knowledge of active venoms having capacity to fully convert the substrate. The inhibitor data was normalised to a percentage inhibition value due to the variation in venom activity.

Statistical analysis of the inhibitors was assessed using two-way ANOVA with Šídák’s multiple comparisons test of each venom control compared to the matched venom treated with inhibitor.

### Bovine plasma coagulation assay

The plasma assay utilises citrated bovine plasma, which is incoagulable until the addition of calcium. Factors involved in the coagulation cascade are targets for specific venom toxins, and so addition of venom and calcium to the citrated bovine plasma can cause the plasma to clot faster than normal (procoagulant toxins) or to not clot at all (anticoagulant toxins). In venom activity experiments, 10 µL of venom at different doses (1 µg – 5 ng) was added to 384-well plates (Greiner, Cat. no. 781101) to determine the most appropriate venom dose for downstream drug inhibition studies. Following a 25-minute incubation at 37°C for consistency with future dose response studies, the plates were acclimatised to room temperature for 5 minutes, before the addition of 20 µL of 20mM CaCl_2_, followed immediately by 20 µL of citrated bovine plasma (Biowest, Cat. no. S0260), which had been centrifuged for 5 minutes at 3000 RCF to pellet any particulate matter. Immediately following plasma addition, the plates were read kinetically on a CLARIOstar Plus at 595 nm absorbance at 25 °C. To provide comparable procoagulant clotting profiles, venom doses were selected that induced complete clotting before the negative control (no venom) wells had initiated clotting. Preliminary drug-only experiments were conducted to identify any inherently pro-or anti-coagulant effects, with only compounds that had no inherent effect on coagulation progressed to dose-response venom inhibition experiments. In these experiments, dose-response curves of inhibitory compounds (as identified in the previous *in vitro* assays) were created in DMSO and 0.5 µL stamped per well of a 384-well flat-bottomed plate at 100x the desired final concentration (final doses of 80 µM – 0.8 nM for all compounds except DMPS tested at 320 µM – 1.3 nM due to reduced potency). Thereafter the experimental workflow was as described above with the addition of venom, preincubation of venom and drug, followed by CaCl_2_ and plasma. As with other assays, AUCs were calculated and normalised to the positive and negative controls to determine percentage inhibition, which was used to plotted as dose response curves, and generate EC_50_ values.

Statistical analysis of the inhibitors was assessed using one-way ANOVA with Dunnett’s multiple comparisons test to the DMSO control for the individual inhibitors and with Šídák’s multiple comparisons for the combined treatments.

### Human blood thromboelastography

The clotting profiles of two representative venoms (*B. atrox* and *B. jararaca*) and the effects of the SVMP and SVSP inhibitors marimastat and nafamostat were tested using thromboelastography ^26,39,57,69–71^. Blood from healthy consenting donors were collected according to ethically approved protocols (LSTM research tissue bank, REC ref.

11/H1002/9) and used up to 4 hours post-sampling with three independent replicates per experimental condition. The blood was collected into BD Vacutainer tubes with ACD-A anticoagulant solution (sodium citrate: 22.0 g/L, dextrose: 24.5 g/L, citric acid: 8.0 g/L and antimycotic [potassium sorbate] reagent: 0.15 g/L (Fisher Scientific, Cat. no. BD 366645)). For each experiment 1.2 mL of whole blood was pre-heated at 37 °C for 5 minutes on a heat block. During this time all reagents were added into the pre-heated sample cup at the following volumes: 12 µL venom sample or PBS control, 15 µL drug sample or PBS control and 20 µL CaCl_2_ (Star-tem, Cat. no. 503-01), before the final addition of 253 µL of whole blood. This resulted in a preincubation time of venom and drug of less than 3 minutes.

Viscoelasticity data were then recorded immediately at 37 °C for 60 min using a ROTEM Delta™ (Werfen). Venom concentrations were at 0.6 µg/reaction (12 µL of 50 µg/mL) diluted in PBS from the 10 mg/mL venom stock. The drug concentrations were diluted in PBS from the 10 mM stock to a 5 µM reaction concentration (15 µL of 100 µM working solution). The negative control (corresponding to spontaneous coagulation of whole blood following recalcification) consisted of no venom or drugs but included CaCl_2_. The positive control consisted of venom and CaCl_2_ without drug treatment. To ensure the CaCl_2_ did not interfere with the drug treatments, each drug was run with CaCl_2_ but no venom. The parameters assessed by ROTEM include coagulation time (CT) and maximum clot firmness (MCF), visualised graphically in an XY plot to demonstrate a clotting profile using Prism v11 software (GraphPad). Additional measures of clot formation time (CFT), alpha angle, and amplitude in 5 minutes (A5) were also reported. These parameters are defined as follows: CT (seconds), time from the start of the measurement until the initiation of clotting classified as 2 mm amplitude; MCF (mm), maximum amplitude of clot firmness reached during the run time, used as a proxy for ‘clotting strength’; CFT (seconds), time interval between the initiation of clotting (2 mm amplitude) until a clot firmness of 20 mm is achieved; alpha angle (◦), the angle between the baseline and tangent to the clotting curve through the 2 mm point. Each parameter is dependent upon different elements of the clotting process. CT represents the coagulation activation via the enzymatic activity of coagulation factors. CFT and alpha angle are dependent on thrombin generation, platelet count/function as well as fibrinogen levels and fibrin polymerization. MCF and A5 are dependent on platelet count/function, fibrin concentration/formation and factor XIII ^26,72^. The outputs from the thromboelastography profiles are calculated by the integrated software on the ROTEM sigma (Werfen). Statistical analysis of the CT and MCF data were analysed independently using one-way ANOVA with Tukey’s multiple comparisons test.

### Chicken egg *in vivo* assay of haemotoxicity

Egg embryos have previously been used to assay venom pathology (e.g., haemorrhage, coagulation, inflammation and lethality) in a vascularised environment and as an efficacy read out for snakebite treatments ^65,67,73^. Fertilised chicken eggs (Medeggs Ltd, UK) on day 1 post-fertilisation were placed horizontally in an incubator at 37°C until day 5, at which point they were sprayed with 70% ethanol and candled to mark embryo position. Damaged and infertile eggs were discarded. A sterile windowing procedure was then conducted to provide visibility of embryos; 6-8 mL albumin was removed using a 23G needle and 10 mL syringe, and the shell was reinforced with clear tape before the marked area was removed using sharp dissection scissors. Eggs were then covered with parafilm and returned to the incubator until day 6. Venom neutralisation studies for marimastat, DMPS or nafamostat were tested at a drug dose of 0.5 µg, 1 µg or 5 µg per egg, resuspended in 5% DMSO, with control experiments demonstrating no effect of 5% DMSO or drug only activity on embryo pathology. On day 6 post-fertilisation, eggs were randomly assigned to dose groups. The group size was 5 for all drug treatment groups, with control groups having increased sample sizes via incorporation across multiple independent experiments (PBS negative control, n=12; venom only control, n=20). Egg embryos were dosed with 10 µL PBS, or 20 µg *B. atrox* venom in 2 µL PBS (5% DMSO) followed by either 9 µL PBS (5% DMSO) or 9 µL treatment (5% DMSO). Doses were pipetted directly onto the vitelline vein on the ventral side of the embryo, with second doses applied immediately after the first at the same location and thus no preincubation of drug with venom. Embryo survival was determined thereafter using observation of the embryo’s heartbeat at multiple timepoints, up to 6 hours, and plotted as Kaplan-Meier survival curves. Pathology was also monitored prior to dosing and throughout the time course using a microscope (Motic SMZ-171) and a mounted smartphone used to capture representative images of selected embryos at a consistent magnification and orientation. At the end of the time course, any surviving embryos were culled by cervical dislocation. Statistical analysis of resulting survival curves was assessed using Log-rank tests (Mantel-Cox) with Holm-Šídák’s multiple comparisons test against treatment vs venom-only control.

## Data analysis

All calculations (including EC_50_ and AUC) and figures (including dose response curves and Kaplan-Meier survival curves) were generated using Prism v11 software (GraphPad). The sample sizes for HTS assays including the SVMP, PLA_2_ and coagulation experiment are the average of the means from independent assays (n >2 within each independent assay). For the serine protease assay, thromboelastography and chicken egg model the sample sizes are individual values due to lower throughput and venom availability.

---

**Supplementary Figure 1.**
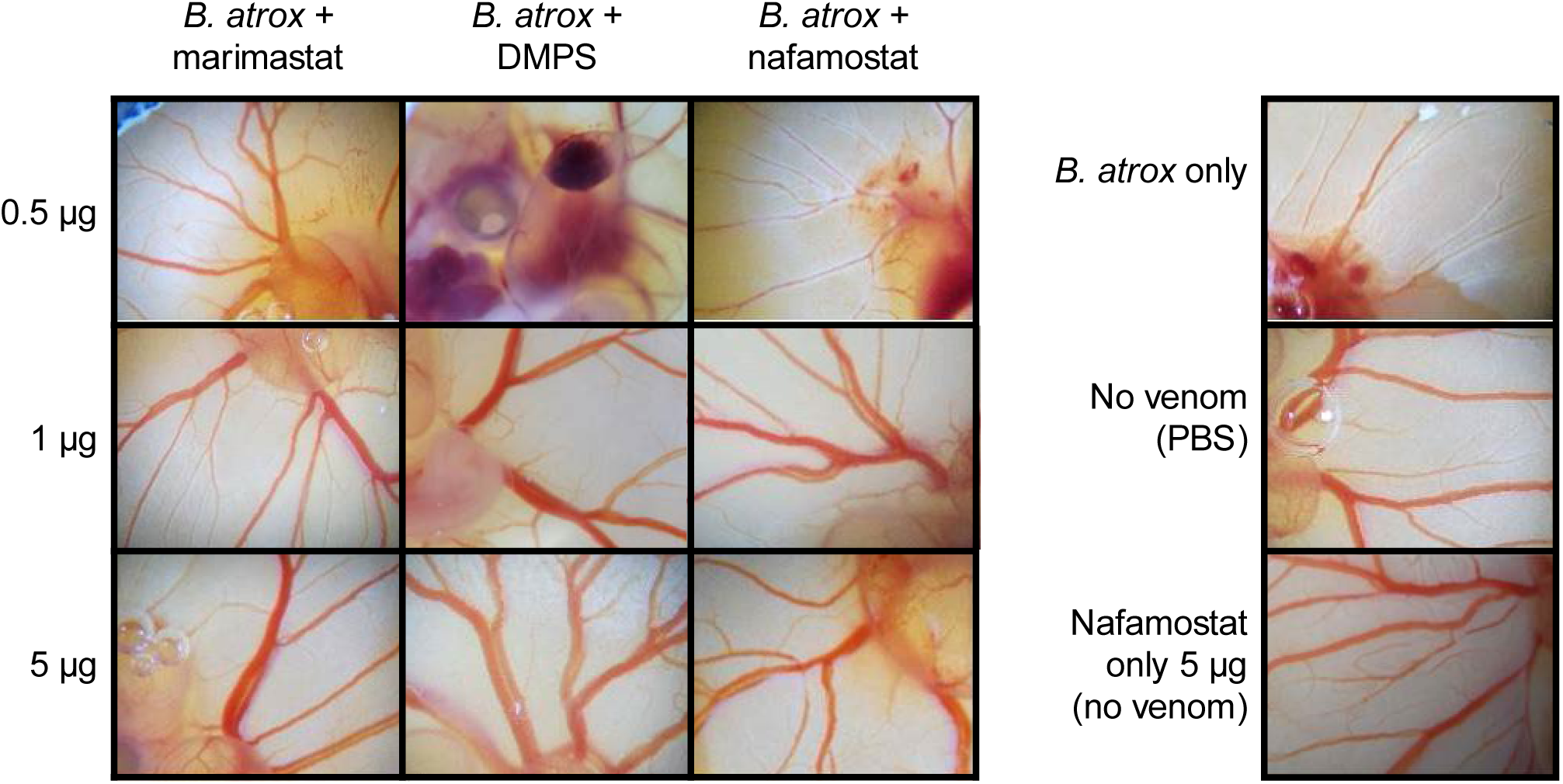
Representative pathology images of chicken embryos 1 hour post-dosing with *B. atrox* venom with and without representative small molecule drugs. Groups of 6-day post-fertilisation chicken egg embryos were dosed on to the vitelline vein with *B. atrox* venom (20 µg) with or without a subsequent dose of 0.5 µg, 1 µg and 5 µg of the inhibitory drugs marimastat, DMPS and nafamostat. The panel on the right includes representative images of control embryos dosed with venom only, no venom or nafamostat only (5 µg).

**Supplementary Table 1.**
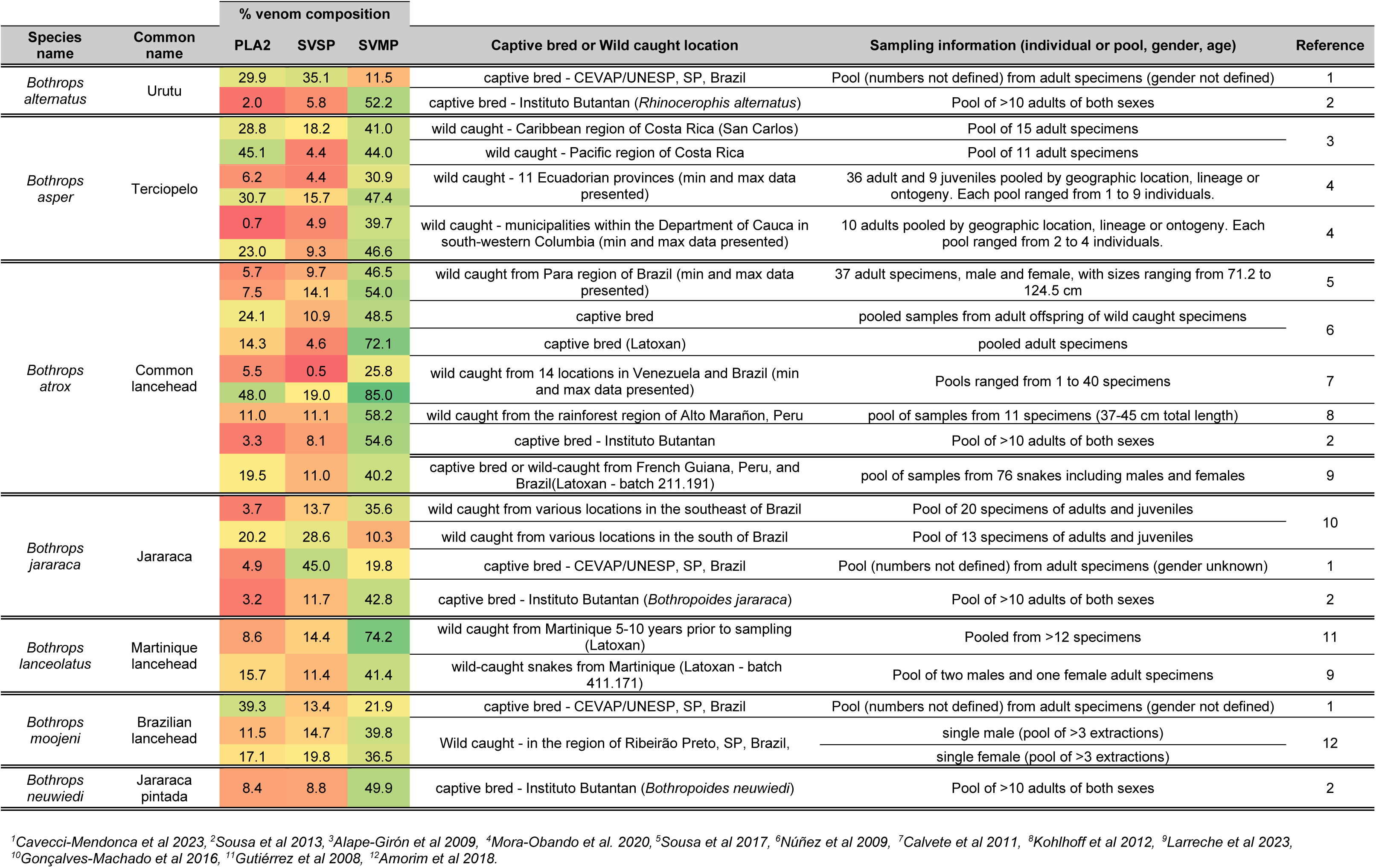
Bothrops Venom Compositions from published proteomics data.

## Notes

### Competing Interest Statement

The authors have declared no competing interest.

### Summary of Updates

Updates have been applied based on the eLife published reviewers comments. These include no additional data or changes to data. Additionally wording to clarify points and additionally statistics have been added.

